# Quantifying unusual neurological movement phenotypes in collective movement phenotypes

**DOI:** 10.1101/2021.07.03.450923

**Authors:** Kehinde Owoeye, Mirco Musolesi, Stephen Hailes

## Abstract

Building models of anomalous behaviour in animals is important for monitoring animal welfare as well as assessing the efficacy of therapeutic interventions in preclinical trials. In this paper, we describe methods that allow for the automatic discrimination of sheep with a genetic mutation that causes Batten disease from an age-matched control group, using GPS movement traces as input. Batten disease is an autosomal recessive lysosomal storage abnormality with symptoms that are likely to affect the way that those with it move and socialise, including loss of vision and dementia. The sheep in this study displayed a full range of symptoms and during the experiment, the sheep were mixed with a large group of younger animals. We used data obtained from bespoke raw data GPS sensors carried by all animals, with a sampling rate of 1 sample/second and a positional accuracy of around 30cm. The distance covered in each ten minute period and, more specifically, outliers in each period, were used as the basis for estimating the abnormal behaviour. Our results show that, despite the variability in the sample, the bulk of the outliers during the period of observation across six days came from the sheep with Batten disease. Our results point towards the potential of using relatively simple movement metrics in identifying the onset of a phenotype in symptomatically similar conditions.

## Introduction

The analysis of the movement patterns of different parts of the body such as eyes [1], muscles [2] and legs [3] in humans [4,5] and animals is important in assessing their overall welfare. Investigating abnormal locomotion patterns, however, is key towards early diagnosis of a number of neurodegenerative diseases such as Batten and Huntington disease. Due to the insidious progression [6] and terminal nature of these diseases, efficient and reliable markers are becoming increasingly important to ensure people with this neurological diseases are given adequate and timely care, attention and treatment and also that the impact of such treatment can be assessed objectively. Batten disease is a rare and fatal autosomal neuro-degenerative disorder [7], characterized by abnormal and involuntary movement patterns (choreoathetosis) [8], personality changes [9], loss of vision [10], and loss of muscle control (ataxia) [11] in its sufferers. It is known to be the most common form of a group of disorders called Neuronal Ceroid Lipofuscinoses (NCLs) and caused by autosomal recessive mutations in the CLN gene [12]. The gradual progression of this disease and consequent worsening of symptoms has a profound negative impact on the quality of life of its sufferers, making them less independent with time and resulting in death in the medium term.

The need to develop therapeutic intervention schemes to be tested has necessitated the use of animals [13] such as mice [14] [15], and sheep [16] for carrying out several studies in this respect. Sheep have brains that are closer to human than are those of rodents, both in terms of size and structure. They also have a longer life-span than rodents hence their choice for this study. Of interest amongst the symptoms associated with sufferers of this disease is the abnormal movement [8] pattern that is usually as a result of gradual changes in the structure of the brain as the disease progresses. This abnormal movement pattern can serve as a biomarker towards identifying the onset of a neurodegenerative disorder such as Batten disease. However, doing this by manual visual inspection can be laborious, time-consuming, inefficient and unreliable (Fig. 1). While using the distance covered in its pure form over the period of observation seems like a plausible option, it turns out that this metric is highly unreliable and uninterpretable (Fig 1). There is therefore a need to define reliable biomarkers for automatic detection and quantification of abnormal movement patterns in this setting.

**Fig 1.**
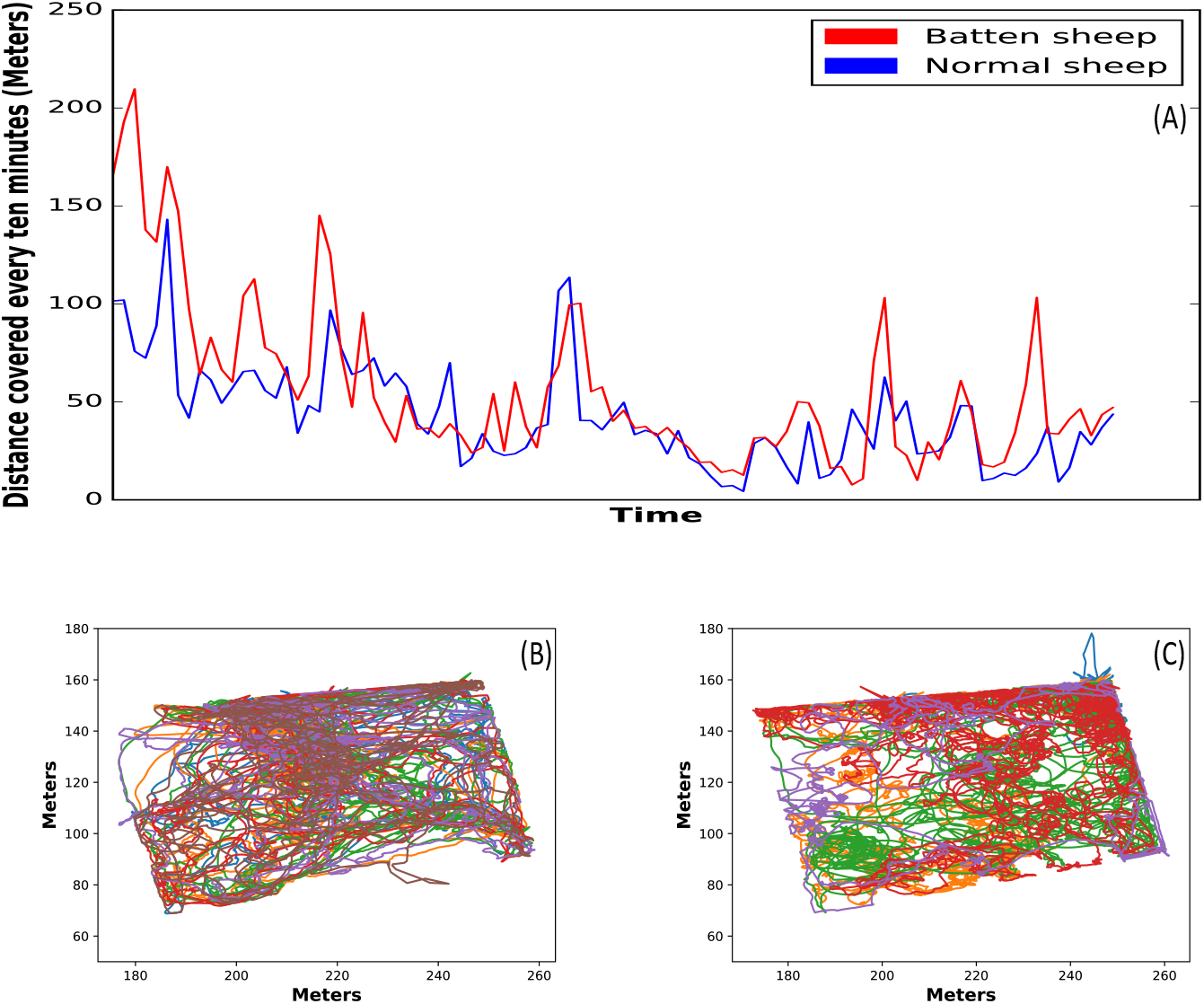
Distance covered and trajectories of the sheep. (A) Mean distance covered every ten minutes by the two groups of sheep on a typical day. (B) Trajectories of the normal/control sheep. (C) Trajectories of the Batten sheep. The mean distance covered by the two groups show its highly unreliable same as visualizing the trajectories towards identifying the sheep with Batten disease.

There has been a significant number of works in the area of automatic methods for welfare of animals using sensor data [17–21]. For example, Beyan & Fisher [18], used a hierarchical classifier to detect novel as well as abnormal fish movement in their aquatic habitat. Using features such as velocity, acceleration and turn angle among others, their corresponding principal components were extracted. The principal components were clustered and together, with the labeled data, a hierarchical classier that leverages similarity of data was built and showed good classification accuracy. Several authors have also proposed different methods to automatically diagnose the health of animals by monitoring their cough using intelligent systems. While Giesert et al. [22] used a Hidden Markov Model to classify coughs into two groups (coughs from healthy pigs and coughs from pigs with respiratory infections), Sa et al. [17] used a motion history image-based method to differentiate the motion peculiar to a coughing animal from other movement patterns. In a similar vein, a neural network was used to analyze patterns in pigs vocalizations in order to distinguish between sick and healthy animals [23]. While with aid of several machine learning algorithms, an early warning system was developed to detect sick broilers in [24]. More recently, a novel method to detect pain levels in sheep was proposed by [25]. They showed it is possible to estimate the pain level in sheep using facial features alone using the Histogram of Oriented Gradients to describe the facial features, followed by training a separate classifier for each feature using a Support Vector Machine. Entropy was also used in [26] to characterize the movement patterns of sheep with Batten disease with their findings revealing Batten sheep on the average have lesser movement entropy compared to the normal sheep. To measure the quality of sleep as well as investigate other forms of neurological dysfunction, Perentos et al. [27] carried out an electroencephalography study of sheep with Batten disease. Their findings reveal that sleep abnormalities as well as epileptic waveforms associated with children with Batten disease were also found in these sheep thus demonstrating the relevance of using these species for *in vivo* studies of human degenerative diseases. No previous work however has tried to address the problem of quantifying abnormal movement phenotypes in collective phenotypes using sheep as subjects.

In this work therefore, we investigate methods for automatically identifying sheep with abnormal movement patterns in a flock. Our approach is one that uses simple movement metrics (distance covered) as a feature to identify outliers within the flock. More specifically, we show that by looking for outliers in the distance covered every ten minutes in the flock and consequently aggregating them, we can with good confidence identify the sheep with Batten disease in a flock. We also show that by finding clusters in the distance covered every ten minutes in a larger flock, we can identify some of the sheep with Batten disease in the cluster with the least membership. In simple terms, our method leverages the herd movement pattern in sheep to classify the sheep whose movement patterns in a certain period deviate from the majority of the flock as abnormal. We evaluate these methods using real world sheep movement data.

## Materials and methods

### Animals

Eleven (5 Batten disease ewes carrying a mutation in CLN5 [28] and 6 healthy ewes age matched control) with mean age of 2 years were maintained at from Lincoln University, New Zealand (NZ) were reared under appropriate procedures approved by the Lincoln University Animal ethics Committee in compliance with the NZ Animal Welfare Act (1999) and in accordance with US National Institutes of Health guidelines. The sheep had water and grazing available to them throughout the period of the study.

### Data acquisition & processing

Movement data for this study were collected for six days using bespoke GPS loggers, details of which can be found in [29]. These were attached to the back of the sheep as shown in Fig. 2. Previous work has shown that such harness equipment carried by sheep does not affect their locomotion [30]. Notwithstanding that sheep are slow moving creatures, we collected GPS data at a sampling rate of 1 sample/s to ensure all forms of interesting movement patterns by the sheep are captured. The data were further processed using the open-source RTKLib and Geographic Library [31] to obtain Cartesian co-ordinates with respect to a local projection. All missing data were interpolated between the last and next seen co-ordinates.

**Fig 2.**
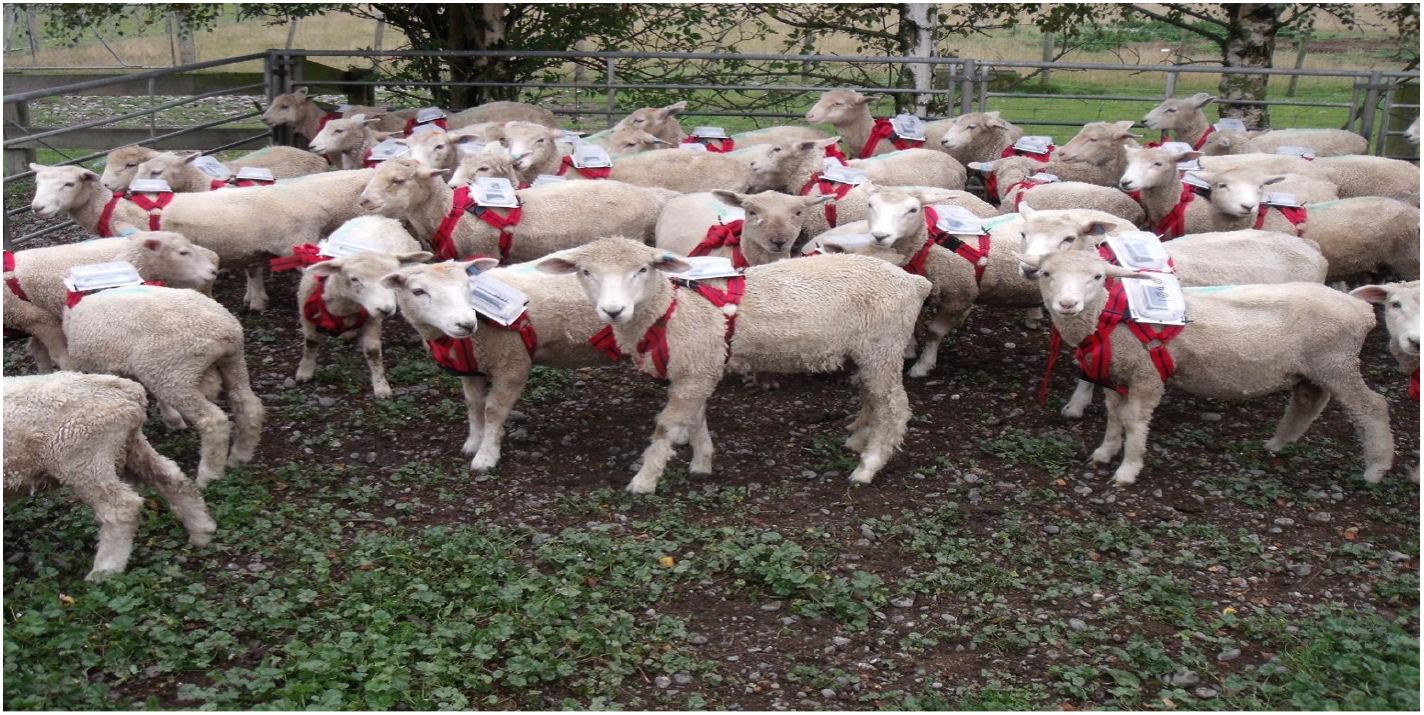
Loggers on sheep. Sheep with GPS loggers attached to their back. Research has shown that such harness equipment does not affect their locomotion [30].

The datasets were all roughly 24 hours in length. Each dataset comprises a phase in which loggers were attached to each sheep in a holding pen; a phase in which the sheep were herded into the field; a phase in which the sheep were left to roam across the field; and a final phase in which the sheep were herded back to the holding pen to have the logging device removed and re-charged. All computations were made based on the data obtained when the sheep were left to wander in the field on their own as this represents the period in which their natural behaviour was most likely to be observed. The eleven sheep were kept together during the first three days of the experiment, and with another group of sixty nine sheep that had also been kept together as well for the remaining three days. The sixty nine sheep consisted of younger animals in a mixture of unaffected sheep and other sheep at different stages of progression with respect to a variety of medical conditions. For our experiments, we extracted the portion of the data where the animals were on the field until at least one of the GPS loggers attached to the sheep failed due to battery exhaustion. All computations were done using Matlab (R2016a).

### Discretizing the trajectory

For each of the sheep in the flock, we obtained the Cartesian trajectories from the GPS traces and partition them into period of ten minutes. We compute the total distance covered on a per second basis and sum it up over each period of ten minutes across the period of observation. Ten minutes was chosen as we consider it a long enough time to observe any significant difference in behaviour.

### Detecting and aggregating outliers

There are several methods to detect outliers in a dataset such as [32] in which a density-based metric was used, [33] in which the k-nearest neighbour concept was used and [34] in which a distance metric was used. We used three fully unsupervised methods discussed below because most of the methods listed above are either not fully unsupervised with requirements for training and test set or they are less parsimonious. The K-nearest neighbour approach, for example, requires the number of nearest neighbours to be specified; which influences the result of our analysis. The local outlier factor approach, in addition to requiring the specification of the number of nearest neighbours to be used also requires a training and test set. The three methods used therefore are Gaussian mixture model (GMM), K-means and gaussian based model. We expand on these methods further below.

### Models

#### Gaussian Mixture Model (GMM)

This is an unsupervised probabilistic model that assumes all data points are generated from a mixture of Gaussian distributions. This approach incorporates uncertainty and assigns each data point to a cluster with some measure of probability. It is assumed with this approach, the cluster with the smallest membership is the anomalous group.

#### K -means

The K-means algorithm is another unsupervised clustering algorithm that finds K groups in data by iteratively assigning each data point to one of K groups based on its distance to a centroid until convergence. While this is a hard assignment, Gaussian mixture model does a soft assignment. As in GMM, the cluster with the least membership is assumed to be the anomalous group.

#### Gaussian Model-Based

This approach assumes data is generated from a Gaussian distribution and that all data that are more than three standard deviations away from the distribution mean are outliers. We examined two instances by using the distance covered every ten minutes as well as the log of this data. This is because the distance covered does not assume a perfect Gaussian distribution.

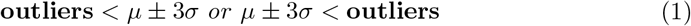

Where *μ* is mean and *σ* the standard deviation. We obtained the outliers from the discretized sub-trajectories for each sheep over the period of observation per day and then aggregate everything over the six days of observation.

## Results & Discussion

### Quantifying abnormal movement behaviour

We used the model and procedures described above, considering only the eleven sheep of interest. We observed that all the sheep with the natural mutation for Batten disease have at least one outlier every day over the period of observation across the four methods whereas this is not the case for the normal (control) sheep group. In addition, it can also be seen that the bulk of the outliers accumulated by the flock over the period of observation across six days belong to the Batten sheep group, as seen in Fig 3, even though they represent a minority (5) of the flock. Also, among the sheep with Batten disease, it can be observed that there is a considerable variation in the total number of outliers accumulated all through the period of observation. This suggests that these sheep have different trajectories in terms of progression of this disease and possible variation in reaction as well. Furthermore, it can be seen that there are no specific patterns in the magnitude of the outliers seen. They can be as low as 10 metres and can be as high as 500 metres using the Gaussian model based (Fig 4), for example, suggesting that these abnormal movements can be random and also a function of how different sheep react to the disease.

**Fig 3.**
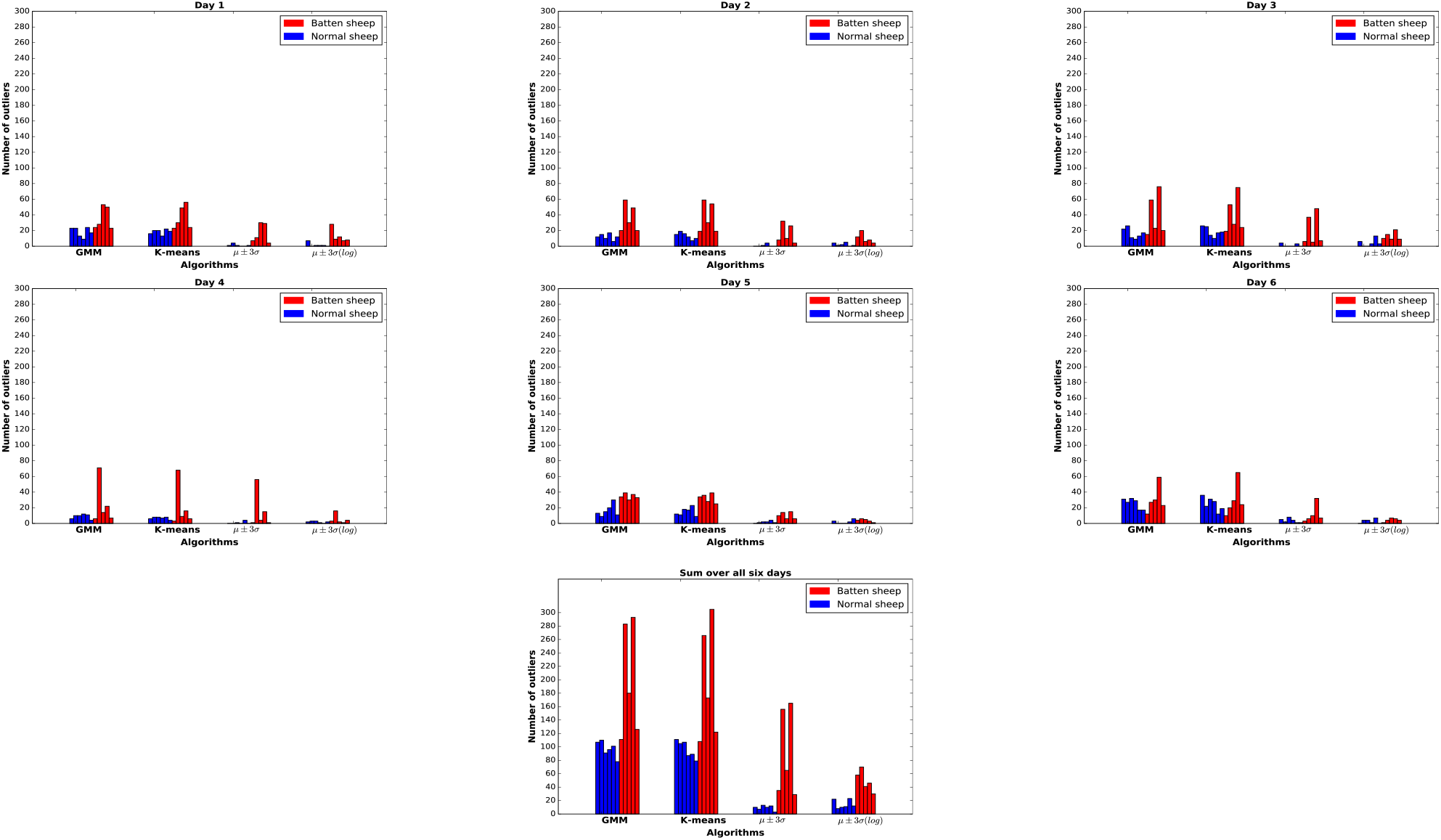
Quantifying abnormal behaviour in a small flock. Bar chart showing the number of outliers detected over six days when the sheep of interest were considered as a flock without others in their environment. It can be seen with all four methods that the sum total of the number of outliers in the batten sheep is greater thus confirming its importance as a biomarker.

**Fig 4.**
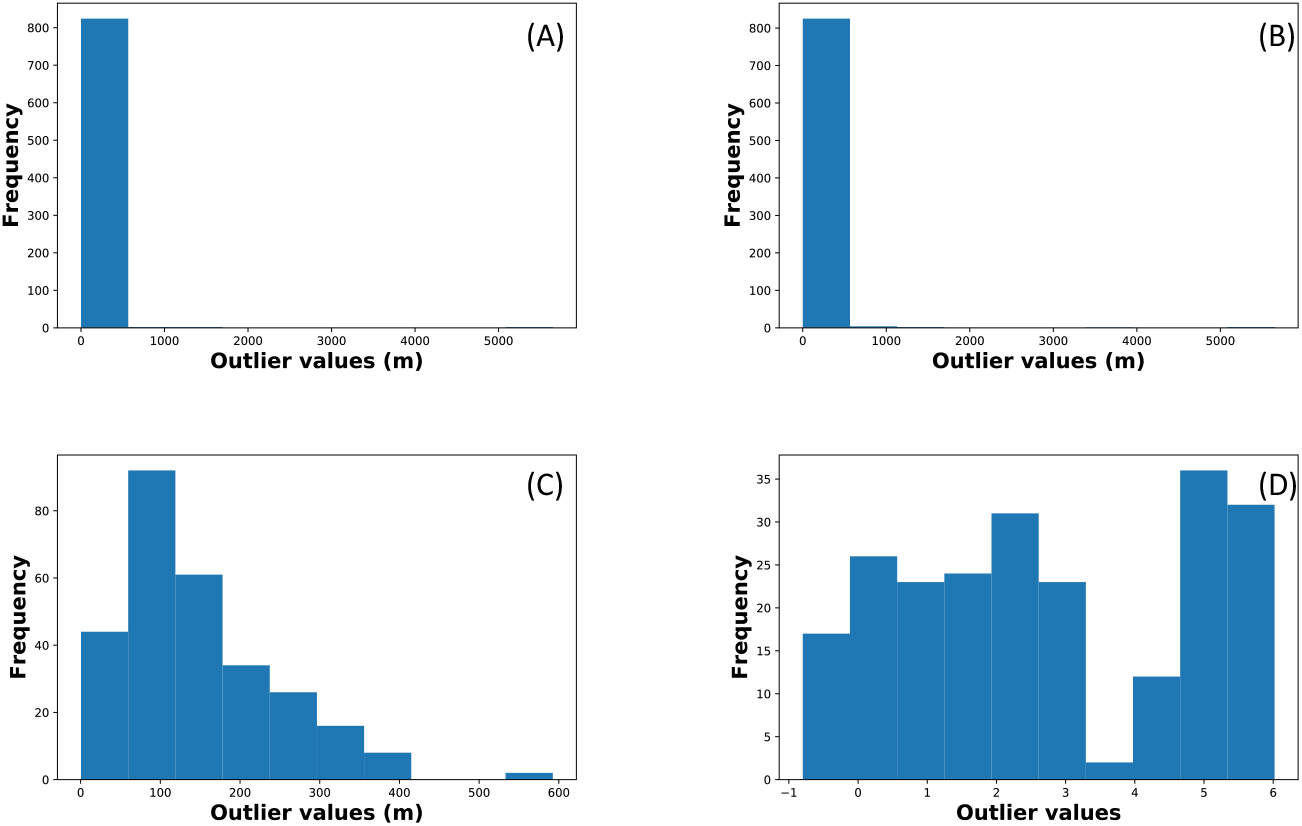
Distribution of outliers. (A) GMM. (B) K-means. (C) *μ* ± 3*σ* (D) *μ* ± 3*σ*(log).

### Quantifying abnormal movement behaviour in a more challenging context

We investigated the possibility of quantifying abnormal behaviour in a more challenging context by considering not only the movement of the Batten sheep relative to their control group but to the whole of the flock during the last three days of the experiment. The previous methods were used initially but results showed they are ineffective because, in a larger group, it is expected that there will be multiple phenotypes in the flock, rather than just two which is the case in this situation. Therefore, we investigated an alternative method. The Akaike information criterion (AIC) was used to select the optimum number between 1 and 20 [35] of groups based on the distance covered every ten minutes in the dataset and then this was used to cluster the sheep into groups using both the Gaussian mixture model as well as the K-means method. We hypothesized that the Batten sheep will most likely be in the group/cluster with the smallest membership. These methods were evaluated on the dataset of movement activities in the last three days of the experiment. Results, show that at least three of the sheep with Batten disease have a conspicuous number of outliers relative to other sheep of interest in this experiment (Fig 5). This can potentially be higher given significant observation period for the experiments.

**Fig 5.**
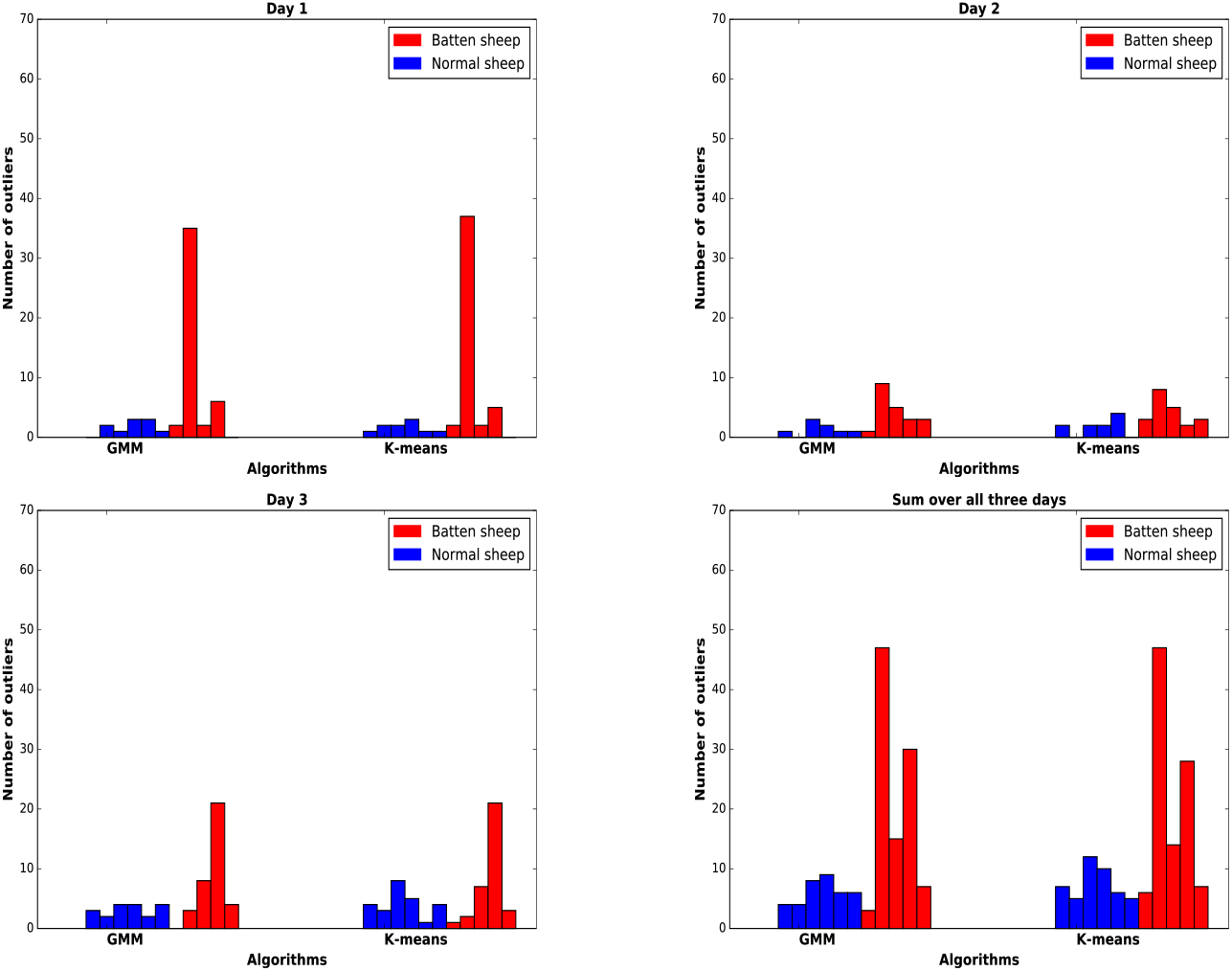
Quantifying abnormal behaviour in a bigger flock. Bar chart of outliers for the two groups of sheep when the movement patterns of other sheep were taken into considerations on the (a) First day, (b) Second day. (c) Third day. (d) The sum over three days. It can be seen that at least three of the Batten sheep have a conspicuously higher number of outliers relative to the rest.

### Statistical analyses and phenotype identification

Learning how significant the difference in the abnormal behaviour of the sheep in this study is important for example, in assessing the efficacy of therapeutic interventions. To further evaluate the performance of all methods, we proposed an equation (2) that computes the ratio of how significant the unusual phenotype quantified here is between the control and the transgenic sheep and how insignificant it is in between the control sheep with respect to the ground truth. Here, it is expected that the pairwise test should be insignificant between the control sheep and significant between the control & transgenic sheep (See example of this in Table 1). We used the Wilcoxon signed-rank test with *p* ≤ 0.05 as threshold to determine significance. Results for the first instance in Table 2 and Fig 6 where the sheep of interest were considered without regard for others in their environment show the difference in behaviour is not statistically significant between the control sheep across the four methods confirming they have a somewhat uniform behaviour (see Table 1 for instance). On the other hand, the difference in behaviour is significant between the two groups of sheep and between the Batten sheep, with the Gaussian model based approach performing best. With respect to evaluation considering other sheep in the flock, we observed no significant difference within the control sheep and some significance between the two groups in Table 2 and Fig 7 with the GMM method performing better than K-means. In addition, the confusion matrices in Figs 6 and 7 show that, while all the methods appear to be adept at recognising the insignificant relationships they struggle in recognising the significant relationships with the Gaussian model based appearing to perform best in this regard in the first scenario and the GMM in the second scenario.

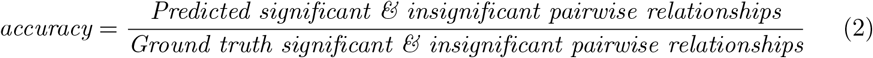

**Table 1.** Pairwise test of significance results using GMM. Where * = Significant when p-value ≤ 0.05, and NS = Not significant when p-value > 0.05. Note: Sheep with IDs 7 to 11 are sheep with Batten disease.

| Sheep ID | Batten Disease | Sheep ID |  |  |  |  |  |  |  |  |  |  |
| --- | --- | --- | --- | --- | --- | --- | --- | --- | --- | --- | --- | --- |
|  |  | 1 | 2 | 3 | 4 | 5 | 6 | 7 | 8 | 9 | 10 | 11 |
| 1 | No | - | NS | NS | NS | NS | NS | NS | NS | NS | * | NS |
| 2 | No | NS | - | NS | NS | NS | NS | NS | NS | NS | * | NS |
| 3 | No | NS | NS | - | NS | NS | NS | NS | NS | NS | * | NS |
| 4 | No | NS | NS | NS | - | NS | NS | NS | NS | * | * | NS |
| 5 | No | NS | NS | NS | NS | - | NS | NS | * | NS | * | NS |
| 6 | No | NS | NS | NS | NS | NS | - | NS | * | * | * | * |

**Table 2.** Significance influenced classification. Accuracy across the two scenarios considered.

| Methods | GMM | K-means | $\mu \pm 3\sigma$ | $\mu \pm 3\sigma$ (log) |
| --- | --- | --- | --- | --- |
| Accuracy - Scenario 1 | 68.33% | 65% | 83.33% | 61.67% |
| Accuracy - Scenario 2 | 66.67% | 60% | - | - |

**Fig 6.**
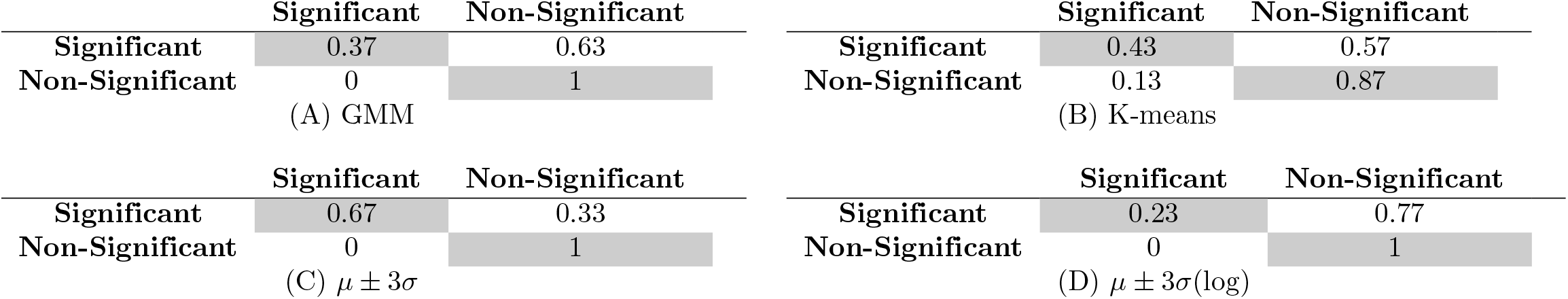
Confusion matrices for the four methods in the first scenario considering the pairwise significant and non-significant relationships.

**Fig 7.**
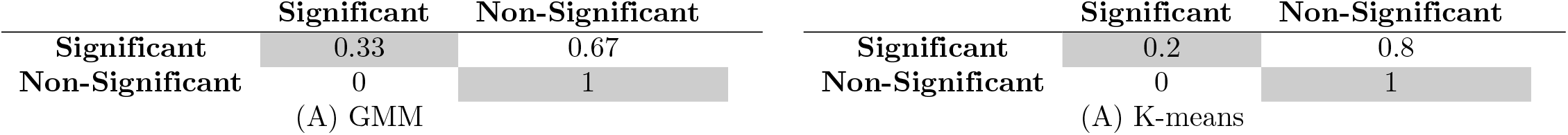
Confusion matrices for the two methods used in the second scenario considering the pairwise significant and non-significant relationships.

## Conclusion

We have investigated and explored multiple methods towards quantifying abnormal movement behaviour and, by extension, identifying sheep with this phenotype in a flock being one of the numerous symptoms associated with sheep with Batten disease. We achieved this by computing the distance covered by every member of the flock within a period whilst looking for outliers in the population. The methods proposed here would not scale in situations where the abnormal sheep are in the majority as the methods are designed for scenarios where the unusual behaviour is in the minority in the population. A class of methods focused on learning collective movement behaviour [36] will be useful here where unusual statistics (behavioural measurements that differs a lot from what is generally known) of the collective behaviour with respect to some known ground truth of normal behaviour can provide useful insights about the welfare of the flock. Whilst the sample size is small during the first three days of the experiment, we devised a method that works well in a challenging environment as experienced in the last three days of the study. The implication of this work transcends identifying sheep with abnormal movement patterns as the methods can be used to monitor the health of individuals or agents that co-exist in groups and have the potential to develop some form of movement disorders during their lifetime. This is a step towards quantifying the progression of a number of neurodegenerative diseases using sheep as subjects. In the future, these methods will be applied to another dataset of transgenic sheep that are expected to display subtle symptoms of Huntington’s disease with the aim of identifying the onset and progression of the condition.

## Acknowledgments

We would like to thank Professor Jenny Morton for key input and helpful discussions. We also acknowledge Professor Alan Wilson, Mr. John Lowe, Dr. Wiebke Scheutt, Dr. Hamed Haddadi of the Royal Veterinary College for their key role in the development of the logging equipment. We thank Professor David Palmer and Ms Nadia Mitchell of Lincoln University for the use of the Batten disease sheep.

## Author Contributions

**Conceptualization:** Kehinde Owoeye, Mirco Musolesi, Stephen Hailes

**Formal analysis:** Kehinde Owoeye

**Funding acquisition:** Kehinde Owoeye, Stephen Hailes

**Investigation:** Kehinde Owoeye

**Methodology:** Kehinde Owoeye

**Project administration:** Stephen Hailes

**Resources:** Kehinde Owoeye, Stephen Hailes

**Software:** Kehinde Owoeye

**Supervision:** Mirco Musolesi, Stephen Hailes

**Writing - original draft:** Kehinde Owoeye

**Writing - review & editing:** Kehinde Owoeye, Mirco Musolesi, Stephen Hailes

